# Ecogenomics of groundwater viruses suggests niche differentiation linked to specific environmental tolerance

**DOI:** 10.1101/2020.07.14.203604

**Authors:** Ankita Kothari, Simon Roux, Hanqiao Zhang, Anatori Prieto, Drishti Soneja, John-Marc Chandonia, Sarah Spencer, Xiaoqin Wu, Sara Altenburg, Matthew W. Fields, Adam M. Deutschbauer, Adam P. Arkin, Eric J. Alm, Romy Chakraborty, Aindrila Mukhopadhyay

## Abstract

Viruses are ubiquitous microbiome components, shaping ecosystems via strain-specific predation, horizontal gene transfer and redistribution of nutrients through host lysis. Viral impacts are important in groundwater ecosystems, where microbes drive many nutrient fluxes and metabolic processes, however little is known about the diversity of viruses in these environments. We analyzed four groundwater plasmidomes and identified 200 viral sequences, which clustered into 41 ~ genus-level viral clusters (equivalent to viral genera) including 9 known and 32 putative new genera. We use publicly available bacterial whole genome sequences (WGS) and WGS from 261 bacterial isolates from this groundwater environment to identify potential viral hosts. We linked 76 of the 200 viral sequences to a range of bacterial phyla, the majority associated with Proteobacteria, followed by Firmicutes, Bacteroidetes and Actinobacteria. The publicly available microbial genome sequences enabled mapping bacterial hosts to a breadth of viral sequences. The WGS of groundwater isolates increased depth of host prediction by allowing identification of hosts at the strain level. The latter included 4 viruses that were almost entirely (>99% query coverage, >99% identity) identified as integrated in the genomes of specific *Pseudomonas, Acidovorax* and *Castellaniella* strains, resulting in very high-confidence host assignments. Lastly, 21 of these viruses encoded putative auxiliary metabolite genes for metal and antibiotic resistance, which might drive their infection cycles and/or provide selective advantage to infected hosts. Exploring the groundwater virome provides a necessary foundation for integration of viruses into ecosystem models where they act as key players in microbial adaption to environmental stress.

**Importance:** To our knowledge, this is the first study to identify the bacteriophage distribution in a groundwater ecosystem shedding light on their prevalence and distribution across metal-contaminated and background sites. Our study is uniquely based on selective sequencing of solely the extrachromosomal elements of a microbiome followed by analysis for viral signatures, thus establishing a more focused approach for phage identifications. Using this method, we detect several novel phage genera along with those previously established. Our approach of using the whole genome sequences of hundreds of bacterial isolates from the same site enabled us to make host assignments with high confidence, several at strain levels. Certain phage-encoded genes suggest they provide an environment-specific selective advantage to their bacterial hosts. Our study lays the foundation for future research on directed phage isolations using specific bacterial host strains to further characterize groundwater phages, their lifecycles, and its effects on groundwater microbiome and biogeochemistry.

## Introduction

Viruses are known to influence the structure and diversity of microbial communities by infection and lysis of microbial cells. Their influence has been widely studied in aquatic communities^1^ where they are predicted to infect approximately one-third of seawater microbes at any given time^2^. In marine ecosystems, major biogeochemical cycles are known to be influenced by viruses affecting community composition, metabolic activity, and evolutionary trajectories^2, 3^. As the recent focus on exploration of viruses in aquatic environments has been on marine ecosystems^4–9^, fresh water environments remained mostly unexplored despite their importance as drinking water supply^10^. The Oak Ridge Field Research Center (ORFRC)^11–13^ is a well-studied United States Department of Energy site that includes groundwater areas with and without metal contamination, referred to as the contaminated and background sites, respectively. It has been well characterized in terms of the physical parameters, microbiome distribution and fluctuation in response to different environmental stresses and thus served as an excellent model groundwater system for studies. We chose this environment to study the incidence of viruses in groundwater microbiome.

Identification of viral sequences in the environment is difficult given the lack of approaches similar to ribosomal DNA profiling in bacteria and their isolation remains challenging because of the difficulties in identifying the bacterial host(s) and our limited ability to culture them. Recently, research has been directed towards exploring viral diversity from metagenome data^7, 14, 15^ thus circumventing these limitations and providing direct insights into the composition of environmental viral communities^16^. In this study we explore an alternate method to sifting through large amounts of chromosomal DNA sequences to find viral sequences by specifically searching circular DNA sequence data generated from the plasmidome analysis. Specifically, we mined the plasmidome data from a well-characterized groundwater system and analyzed the resulting viral sequences complete with genomic and ecological contexts.

## Methods

### 2.1 Groundwater sample collection and sequencing analysis

The groundwater samples were obtained from Oak Ridge Field Research Center (ORFRC) site^11–13^ and included metal-contaminated (wells FW104 and FW106) and background (wells GW456 and GW460) areas. An earlier study^17^ described the circular DNA isolation (plasmidome analysis) procedure from 4 liters of groundwater from background sites (GW456 and GW460), followed by sequencing, assembly, annotation and other analyses. Additionally, for this present study, we also use plasmidome sequence data from two contaminated site samples comprised of 8 liters ground water from FW104 and FW106 and subjected to the same analysis (manuscript in preparation, sequencing data available via MG-RAST IDs mgm4830571.3 and mgm4830867.3).

### 2.2 Identification of viral contigs

Post sequencing, the assembly of all contigs (including plasmid and viral DNA), along with prediction of circular sequences using bioinformatic analyses were performed as described previously^17^. Briefly, all plasmid sequences obtained were subjected to a pipeline method for postassembly detection of circularity among scaffolds, and any scaffolds failing this are termed as non-circular contigs, to distinguish them from those plasmid sequences which passed the criteria. All circular contigs along with non-circular contigs encoding more than 10 proteins were subjected to VirSorter analysis^18^, an iVirus tool available via Cyverse^19^ for identification of viruses. VirSorter was used to identify and remove microbial contigs using the ‘virome decontamination’ mode, with every contig that was not identified as viral considered to be a microbial contig. The final set of viral contigs was formed by compiling sequences detected as VirSorter categories 1 and 2 along with prophage categories 4 and 5 (Table S1). Thus, we focus on the 200 viral sequences with high confidence assignments (VirSorter categories 1,2 4, and 5), and ignored the low confidence assignments (VirSorter categories 3 and 6). Vcontact 2^20^ was used to perform viral cluster analysis, and the results were visualized using cytoscape^21^. Since the groundwater from background site was spiked with strains *Desulfovibrio vulgaris* Hildenborough (ATCC 29579), *Escherichia coli* DH1 (ATCC 33849), and *E. coli* strain J-2561 as controls for the plasmidome study^17^, any viruses associated with these strains were removed from the analysis. Given that the DNA isolation procedure concentrated on targeted isolation of circular DNA, there is an expected inherent bias in identifying circular dsDNA viral sequences from this dataset.

### 2.2 Generation of host database

#### 2.2.1 Generation of host database from ORFRC bacterial isolates

##### Isolation of bacterial strains

The bacterial isolates were obtained via direct-plating under aerobic or anaerobic conditions at 25-30 °C in the dark, using ORFRC groundwater or sediment extract as inoculum, or via two-step isolation: enrichment incubation of 1 ml groundwater in 9 ml liquid media aerobically for two weeks followed by direct-plating for isolation. A subset of isolates were obtained from biofilm reactors (CDC reactors) that were fed ORFRC groundwater and had non-porous glass beads (30 um) as matrix for biofilms in coupons. Water or beads from the reactors were used as inoculum. For direct-plating, rich media (Luria-Bertani, tryptic soy, R2A, Eugon, Winogradsky) agar plates, or basal medium (4.67 mM ammonium chloride, 30 mM sodium phosphate, with vitamin and mineral mixes as previously described^22^) agar plates were used. The liquid media for enrichment incubation was filtered groundwater amended with one or a combination of the following carbon sources: glucose (5 mM), acetate (5 mM), benzoate (0.5 mM), casamino acid (10 μg/ml), bacterial cell lysate, and sediment-extracted dissolved organic matter. After direct-plating, single colonies were picked and regrown in liquid media for 16-48 h until the culture reached mid-log phase. Then a portion of the culture was used to extract DNA for 16S rRNA based identification, and the rest were cryopreserved with sterile glycerol (to a final concentration of 30%), flash frozen with liquid nitrogen, and stored at −80 °C.

##### Whole genome sequencing and de novo assembly

Cultures were revived from glycerol stocks by streaking onto Luria-Bertani or R2A agar plates. Individual colonies developed at 30 °C over 48 h, which were then inoculated into corresponding liquid media and grown at 30 °C for 48 hours. The cultures were centrifuged, the genomic DNA was extracted using the Qiagen DNeasy kit (Qiagen, Venlo, NL) according to the manufacturer’s instructions. All samples were eluted in Qiagen’s AE buffer: 10 mM Tris-Cl, 0.5 mM EDTA, pH 9.0. Genomic DNA was stored at −20 °C followed by transfer into a 384-well plate for automated library preparation. The isolated genomic DNA was normalized to 0.2 ng/uL in 10 mM Tris (pH 8.0), and libraries were prepared using the Illumina Nextera XT kit at 1/12th reaction size on a SPT Labtech Mosquito HV. Final libraries were purified using Solid Phase Reversible Immobilization beads, and sequenced on an Illumina NextSeq 500 with 150 bp paired-end reads. The program Cutadapt v1.12 was used to remove adapter sequences with parameters -a CTGTCTCTTAT -A CTGTCTCTTAT^23^. We performed sliding window quality filtering with Trimmomatic v0.36 (parameters-phred33 LEADING:3 TRAILING:3 SLIDINGWINDOW:5:20 MINLEN:50)^24^. All genomes were assembled de novo using SPAdes v3.9.0 with the following options (-k 21,33,55,77 --careful)^25^. Genome quality was validated with the program checkM v1.0.6 using the lineage_wf pipeline with default parameters^22^, and all draft genomes passed the criteria of contamination < 10% and completeness > 95%. The 16S rRNA gene sequences were recovered with RNAmmer v1.2 (‒S bac ‒m ssu) and taxonomically classified with SINTAX (usearch v9.2.64) against the Ribosomal Database Project (RDP)^28^ 16S rRNA gene training set v16 with species names and the following parameters (–strand both–sintax_cutoff 0.8)^26, 27^. The whole genome sequences (WGS) of 261 bacterial isolates (details in Table S2) from ORFRC were combined to form a database for further bioinformatic analyses. The WGS of the 261 strains are available via (https://kbase.us/n/63776/35) and the DOI (10.25982/63776.53/1637360).

#### 2.2.2 Generation of host database from NCBI bacterial and archaeal isolates

A genome database of putative hosts for the viruses was generated including all archaeal (311 assembled complete genomes, downloaded in September 2019) and bacterial (14028 assembled complete genomes, downloaded in August 2019) genomes from NCBI Assembly. The taxonomic affiliation of the genomes was taken from the NCBI taxonomy.

### 2.3 Host prediction and diversity

Three different previously published approaches^29, 30^ for predicting hosts based on examining similarities between a) bacterial genome encoded CRISPR spacer and viral genome^31^ b) viral and microbial genomes due to integrated prophages or gene transfers^32^ and c) viral and host genome nucleotide signatures (here, tetranucleotide frequency similarity)^33^ were used as described below. The confidence in assignment via these three methods to different clades in bacterial classification has been previously estimated^34^ with CRISPR-based predictions being the most accurate while the tetranucleotide frequency-based predictions were the least accurate at the genus level.

#### BLAST-based identification of sequence similarity between viral contigs and host genome

All 200 viral contigs were compared to all archaeal and bacterial genomes with BLASTn (threshold of 50 for bit score and 0.001 for *E*-value), to identify regions of similarity between a viral contig and a microbial genome, indicative of a prophage integration or horizontal gene transfer. As previously established^30^, host prediction was made when an NCBI genome displayed a region similar to the viral contig ≥4.9 kb at ≥70% identity. When one viral sequence had hits to multiple bacterial strains, the top 5 hits (based on bit score) were analyzed to determine the last common ancestor clade. This clade was then assigned as the host to the virus. Based on this methodology, genus level bacterial host predictions were made. Bacterial strain-specific host predictions were only made when the entire virus was found to be encoded in the bacterial whole genome sequence. In this case, BLAST with highly stringent parameters, referred to as BLAST99 (>99% query coverage, e-value=0 and >99% identity) was performed to query for the presence of an entire viral sequence in the host.

#### Matches between viral contigs and CRISPR spacers

CRISPR arrays were predicted for all ORFRC microbial genomes with CRISPR Recognition Tool, CRT^35^ using default settings (repeat settings used 3 minimum repeats, 19 minimum repeat length, 38 maximum repeat length, and a search window of 8; along with spacer settings used 19 minimum spacer length and 48 maximum spacer length). We used previously published^30, 36^ BLAST parameters for identifying the target of CRISPR spacers (i.e. using the BLASTn-short task, a maximum expect value of 1; a gap opening penalty 10; a gap extension penalty 2; a word size 7; and dust filtering turned off). Given that the accuracy of this approach for detecting phage hosts strongly depends on the maximum number of mismatches allowed between the CRISPR spacer and the viral sequence, the results were filtered to allow 0 or 1 mismatch. Only the CRISPR spacers that matched viral sequences were then compared back with the bacterial WGS with no mismatch to come up with bacterial host predictions. Based on this methodology, strain level bacterial host predictions were made.

#### Nucleotide composition similarity: comparison of tetranucleotide frequency

Bacterial and archaeal viruses tend to have a genome composition close to the genome composition of their host, a signal that can be used to predict viral–host pairs^30, 33, 37^. Here, canonical tetranucleotide frequencies (also referred to as 4mer) were observed for all viral and host sequences using Jellyfish^38^ and mean absolute error (that is, the average of absolute differences) between tetranucleotide-frequency vectors were computed with in-house Perl and Python scripts for each pair of viral and host sequence as previously reported^34^. A viral contig was then assigned if the average of absolute differences (*d*) between tetranucleotide-frequency vectors *d* < 0.001. When multiple strains had hits to one viral sequence, the top five hits (based on lowest distance) were analyzed to determine the lowest common ancestor to the group. This lowest common ancestor was then assigned as the host to the virus. Based on this methodology, genus level bacterial host predictions were made.

### 2.3 Phylogenetic tree construction

For constructing the phylogenetic tree using ORFRC isolates, the 16S rRNA sequences from all 261 strains were aligned using Muscle^39^. The evolutionary history was inferred by using the Maximum Likelihood method based on the Tamura-Nei model using MEGA7^40^. The tree with the highest log likelihood (−7846.44) is shown. Initial tree(s) for the heuristic search were obtained automatically by applying Neighbor-Join and BioNJ algorithms to a matrix of pairwise distances estimated using the Maximum Composite Likelihood (MCL) approach, and then selecting the topology with superior log likelihood value. The analysis involved 255 nucleotide sequences. All positions containing gaps and missing data were eliminated. There were a total of 519 positions in the final dataset. All branches were collapsed at the genus level. For the phylogenetic tree depicting NCBI isolates, existing trees were downloaded using NCBI taxonomy and collapsed to genus levels.

## 3. Viral sequence annotation

A functional annotation of all virus-encoded predicted proteins was based on a comparison to the Pfam domain database v.32^41^ with HmmScan^42^ (threshold of 30 for bit score and 10^−3^ for *E*-value). The Pfam categories were assigned based on Pfam target name as previously described^34^, and any Pfam target name not categorized earlier is referred to as “not categorized”. All contigs were also uploaded to KBase for annotation. To specifically identify metal and antibiotic resistance genes, all the unique Pfam target names and their descriptions were manually curated.

## Results and Discussion

### New viruses detected in the circular DNA datasets

To study groundwater viruses, we leveraged existing data focused on extrachromosomal circular DNA templates by identifying viruses from plasmidome datasets (Fig 1). Viruses and plasmids can coexist stably, support the transfer of each other to new hosts^43^ or even form a hybrid^44^. Given that both can be found as extrachromosomal circular DNA molecules, we used VirSorter, a tool designed to predict bacterial and archaeal virus sequences on the plasmidome assemblies^18^ and identified 200 sequences as groundwater viral sequences from 13,770 plasmidome contigs (Fig S1). We then categorized viral sequences into viral clusters (approximately equivalent to known viral genera) using shared gene-content information and network analytics^33, 45^. Clustering of the 200 groundwater viral sequences with publicly available bacterial and archaeal viruses revealed that 85 groundwater viral genomes formed 41 viral clusters with at least one representative of groundwater virus (Table S3). Of these 41 clusters, 9 included a reference viral genome (Fig 2) and 32 were putative new viral genera. The details on the size of different clusters is depicted in Fig S2. The largest identified virus was a circular 296,356 bp contig (see virus size distribution depicted in Fig 3), and was part of a novel viral cluster. Although more viral sequences were identified from the background versus contaminated groundwaters, the fractions of all contigs identified as viral sequence was similar across both sites (Fig S1). Thus, the 200 groundwater viruses spanned a wide variety of sizes and included representatives of both known and novel viral genera.

**Fig 1:**
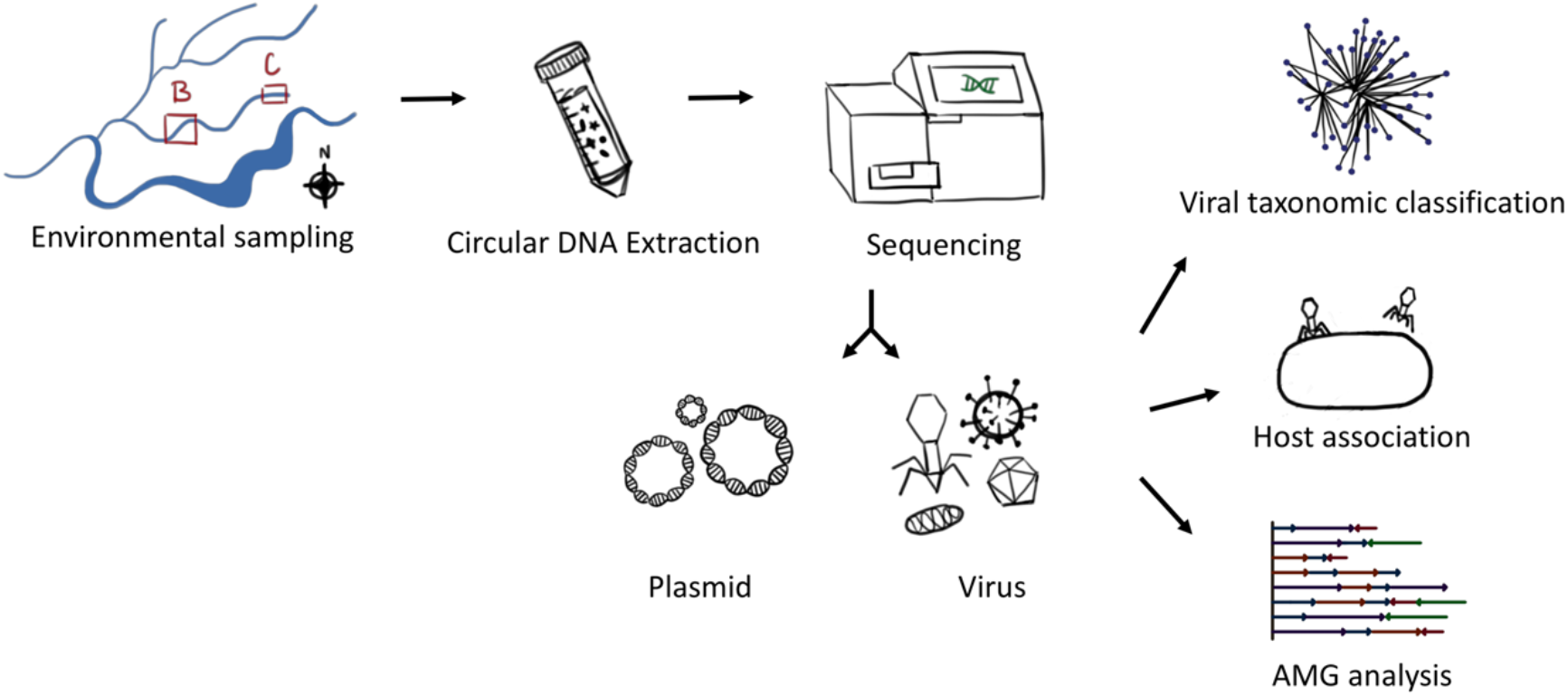
Overview of the study. Groundwater from the Oak Ridge Field research Site from background (B) and contaminated (C) areas was filtered and subjected to circular DNA extraction. Sequencing, assembly and annotation resulted in identification of both plasmids and viral genomes. The viral genomes were subjected to viral cluster analysis to study the virus types, host association analysis to get a prediction of bacteria they might infect and Auxiliary Metabolite Analysis (AMG) analysis to study what functional genes they encode.

**Fig 2:**
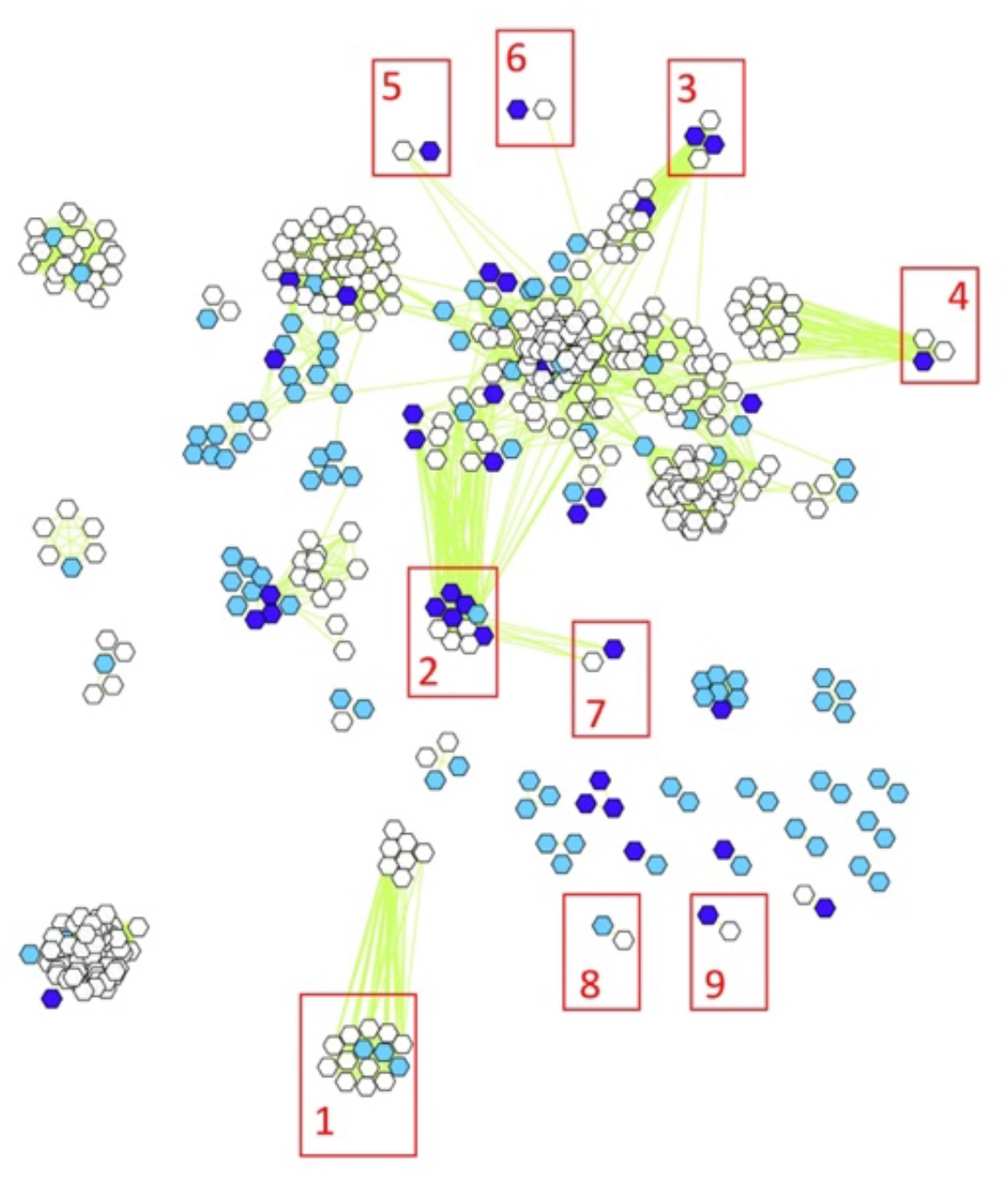
vContact generated viral cluster map depicting clustering of 85 viral sequences from background (light blue) and contaminated (dark blue) groundwater, along with known virus reference genomes (white). The 9 viral clusters that contain known viruses are annotated on the figure as 1) *Microviridae* 2) *Podoviridae* (*Caudovirales*) 3) *Myoviridae* (*Caudovirales*) 4) *Myoviridae* (*Caudovirales*) 5) *Podoviridae* (*Caudovirales*) 6) *Siphoviridae* (*Caudovirales*) 7) *Podoviridae* (*Caudovirales*) 8) *Inoviridae* and 9) *Myoviridae* (*Caudovirales*). The order and distance between different viruses is an arbitrarily selected value.

**Fig 3:**
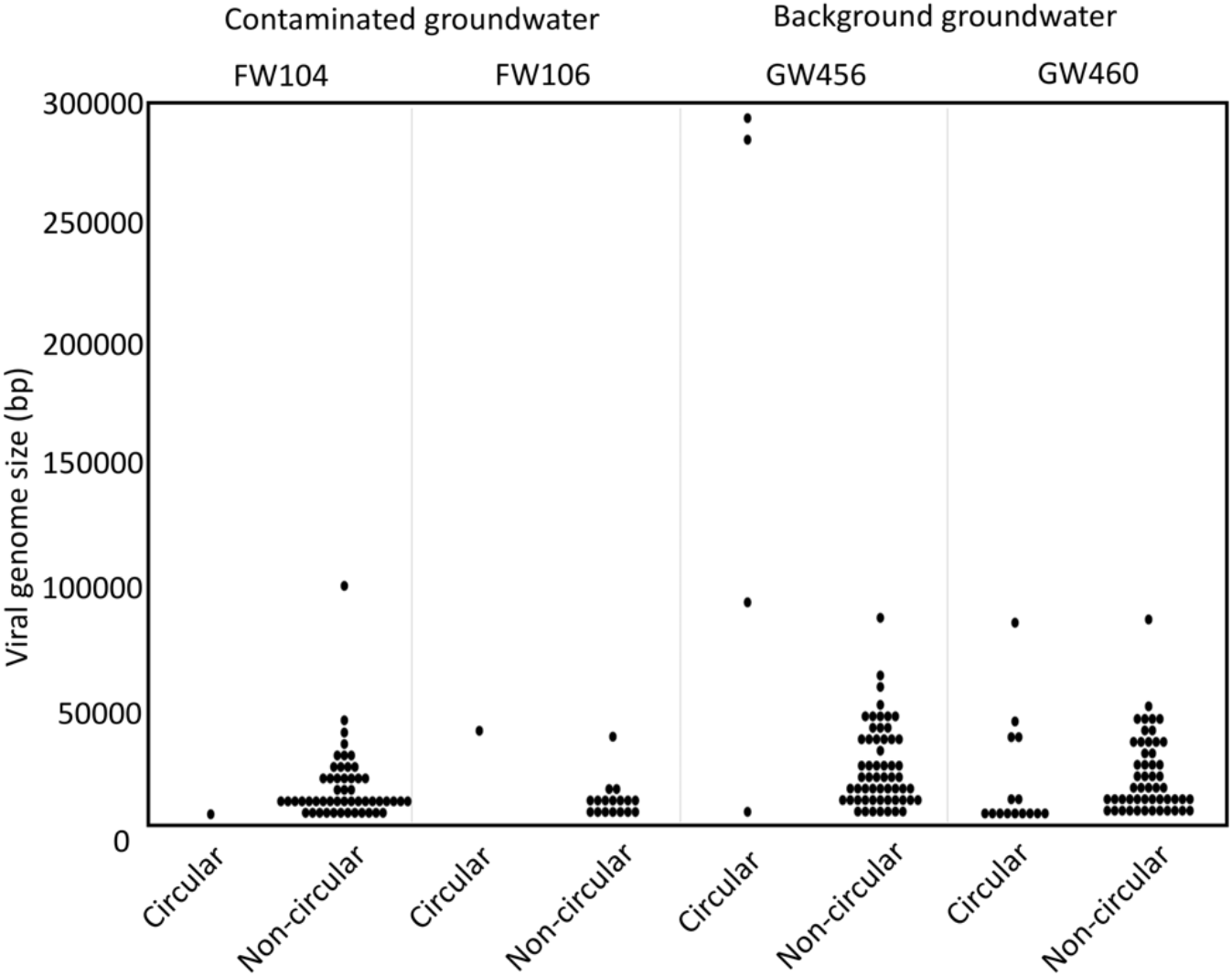
Size distribution of viruses from the background and contaminated groundwaters.

Several aspects of the viral clusters provide evidence to optimal clustering of groundwater viruses. All the 9 viral clusters with known reference viral genomes were circular DNA viruses. The VC with 14 representatives had 11 representatives belonging to the family *Microviridae*, sub family *Gokushovirinae*, which are 4.5–6kb, circular single stranded DNA virus. Interestingly, the 3 viral contigs that are clustered are from the background site, and are also in the same size range (4.61, 4.78 and 5.09 kb). At least one virus (GW460_nc_scaffold_3616, 8250bp size) from background site is an inovirus (5-15kb size, circular single-stranded DNA genomes with rod-shaped or filamentous virions^46^) clustering with known inovirus *Ralstonia*~phage~1~NP-2014. The genome of inoviruses are known to be chromosomally integrated or replicated as a plasmid^47^, which may be why this virus was recovered from plasmidome data.

### Host Predictions

Once we identified viral genomes and their clusters, we sought to identify the range of hosts that these viruses infect. Using the 261 ORFRC bacterial isolates we were able to assign bacterial hosts to 20 viral genomes (Fig 4) out of the 200, indicating we were able to predict hosts for 10% of the viral genomes identified (Table S4). As expected, the maximum number of predictions were made using tetranucleotide frequency (16), followed by BLAST (9) and CRISPR (2) analysis (Fig S3). All 9 viral sequences that had bacterial host genus predicted via BLAST, also had strain level predictions using BLAST99. An example of host prediction via BLAST99 is depicted in Fig S4 where the entire viral sequence was found in five different *Acidovorax* strains. Interestingly, 7 viral genomes were assigned hosts using both BLAST and tetranucleotide frequency methods and 6 of them were predicted to the exact same bacterial genus, increasing the confidence in their host prediction. Out of 20, 10 viral genomes had *Pseudomonas* predicted as its bacterial host, and overall 18 viral genomes were assigned to Proteobacteria. This could be attributed to the fact that out of 261 ORFRC isolates, over 50% were *Pseudomonads*, and over 85% were Proteobacteria, making it easier to identify them as host strains. Thus, several ORFRC bacterial genus and strains belonging to phyla Proteobacteria, Actinobacteria, and Firmicutes were predicted as hosts for the viruses.

**Fig 4:**
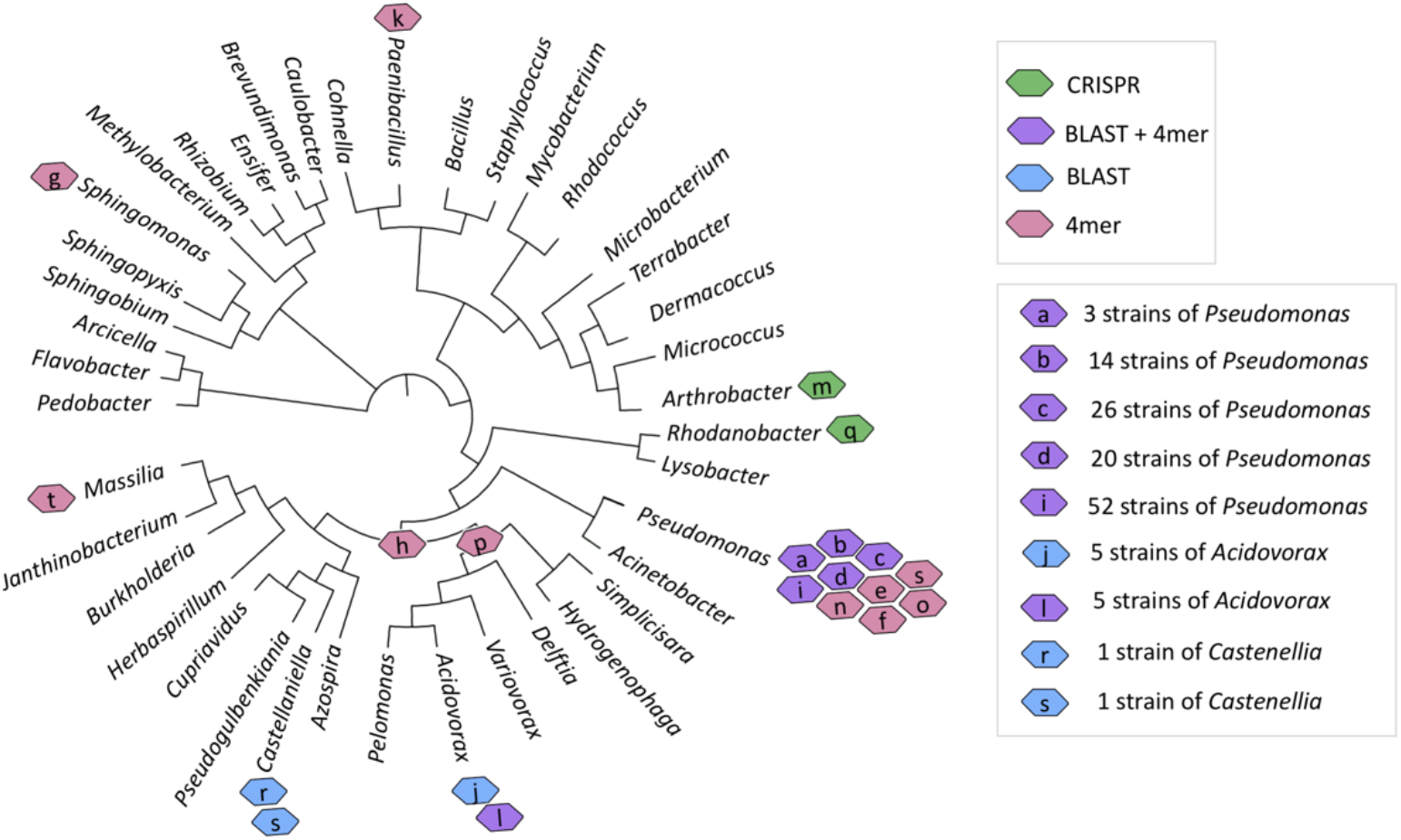
Viral host predictions based on BLAST, high stringency BLAST (BLAST99), tetranucleotide frequency (4mer) and CRISPR methods using whole genome sequence(WGS) information from 261 ORFRC bacterial isolates. The details of the 20 viruses (“a”-“t”) are provided in Table S4. The viruses “h”, and “p” have their hosts assigned to Class Betaproteobacteria and Family Comamonadaceae. The rest of the viruses are assigned to the genera. The phylogenetic tree was made from 16S rRNA sequence of 261 ORFRC isolate strains. The viral sequence “s” appears twice because it was predicted to infect two different genera based on the different prediction methods.

We also leveraged the complete archaeal and bacterial genome sequences available on NCBI, to make predictions of bacterial hosts for the 200 viral genomes. No hits were found using the 311 archaeal strains. Using the 14,028 bacterial strains, host predictions could be made for about 36.5 % (73 out of 200) of the viral genomes, with a vast majority assigned to the phylum Proteobacteria (Table S5). Other bacterial hosts were in the phyla Actinobacteria, Bacteroidetes, Firmicutes, Chlamydiae and Chloroflexi. Again, the maximum number of predictions were made using tetranucleotide frequency (71), followed by BLAST (5) analysis (Fig S3). The BLAST99 had no hits, so strain specific bacterial host predictions were not made. Interestingly all 5 viral genomes that had predictions with BLAST, also had predictions using tetranucleotide frequency method. Although a higher number of viral sequences could be assigned to bacterial hosts using WGS from NCBI compared to ORFRC, the probability of finding a host for every bacterial WGS tested was higher with ORFRC strains (7.6%) compared to NCBI strains (0.5%), highlighting the benefits of including bacterial strains from the same environment as the viral sequence itself. More importantly, strain-specific host assignments could only be made using groundwater bacterial isolates, and such high-resolution host assignment is important when designing experiments aimed at isolating specific phages.

Together using the ORFRC and NCBI strains host predictions we were able to assign bacterial hosts to 38% (76 out of 200) of the viral genomes (Fig 5). Around 17 viruses had host predictions based on both ORFRC and NCBI strains (Fig S3), with the same bacterial phyla predicted as hosts (Table S6). Differences like this could be attributed to the non-overlapping nature of the strains from NCBI and ORFRC, and differences in the strength of host prediction methodologies. Next, we compared host prediction between members of the same viral cluster (Table S7). The bacterial host predicted mostly remained consistent within the same viral cluster. The minor discrepancy seen in the viral clusters can likely be explained on further analysis, for instance the exceptional viral cluster (VC_138_0), consists of ten members with six being groundwater viruses and their hosts were predicted to be either *Burkholderiales* or *Pseudomonadales* based on the prediction method used. Interestingly, the four known viruses they cluster with were *Bordetella* virus BPP1, *Pseudomonas* phage AF, *Pseudomonas* phage vB_PaeP_Tr60_Ab31, and *Xanthomonas citri* phage CP2 indicating members of this cluster infect both *Burkholderiales* and *Pseudomonadales*. Thus, consistent patterns of host prediction emerge within the same viral cluster.

**Fig 5:**
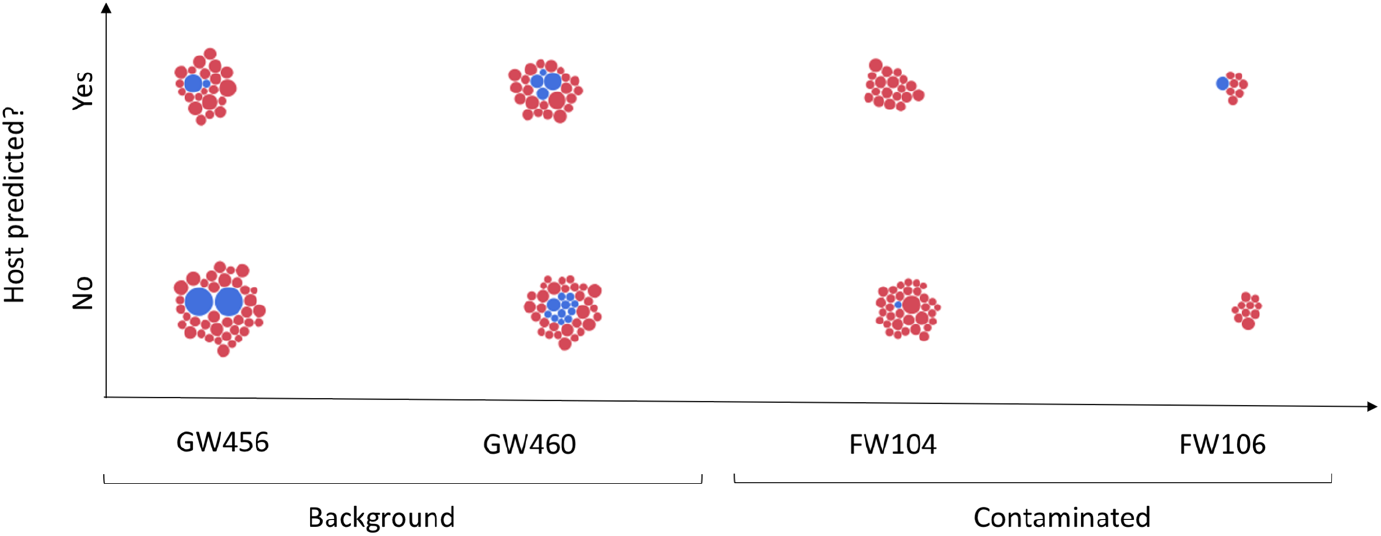
Compilation of viral sequences from the groundwater sites based on availability of bacterial host prediction. Circular viral sequences are depicted in blue, while the rest are in red. The size of the circle is indicative of the viral sequence size.

### Presence of metabolic genes

In addition to affecting groundwater biogeochemistry through their physical contribution to dissolved organic matter and the lysis of their hosts, viruses can also affect the diversity and function of microbial populations through the incorporation and expression of Auxiliary Metabolic Genes (AMGs)^4, 48^. AMG definitions are still being refined^49^, but generally these genes are not involved in viral replication or structure but instead allow viruses to directly manipulate host metabolism during infection. Examination of all the viral sequences revealed a total of 1,486 hits classified into known Pfam categories^34^ (Fig S5). Exploring Pfam domains associated with microbial metabolism resulted in the identification of 51 unique putative AMGs (Table S8). Since these viral sequences are from a site where metal and antibiotic resistance genes are routinely seen^17, 50, 51^, all the unique PFAM hits were manually curated to identify metal and antibiotic resistance genes. We found that the metal resistance genes identified as putative AMGs were those providing resistance to copper, while the antibiotic resistance genes in the list of putative AMGs were annotated as beta lactamase multi-resistance providing resistance to β-lactam antibiotics, multi-drug efflux pumps AcrB/AcrD/AcrF family providing multi-drug resistance, and streptomycin adenylyltransferase providing resistance to streptomycin. An excellent example is viral sequence GW456_c_scaffold_130 which was annotated to encode metal and antibiotic resistance genes along with signature phage genes consistent with a complete phage genome (Fig 6, annotation details in Table S9). The compilation of all the data discussed is available in Table S10. To the best of our knowledge, this is the first report of the presence of metal and antibiotic resistance genes on viral sequences. The presence of metal and antibiotic resistance genes suggests that groundwater viruses may manipulate metal tolerance mechanisms enabling their hosts to adapt to environmental stressors.

**Fig 6:**
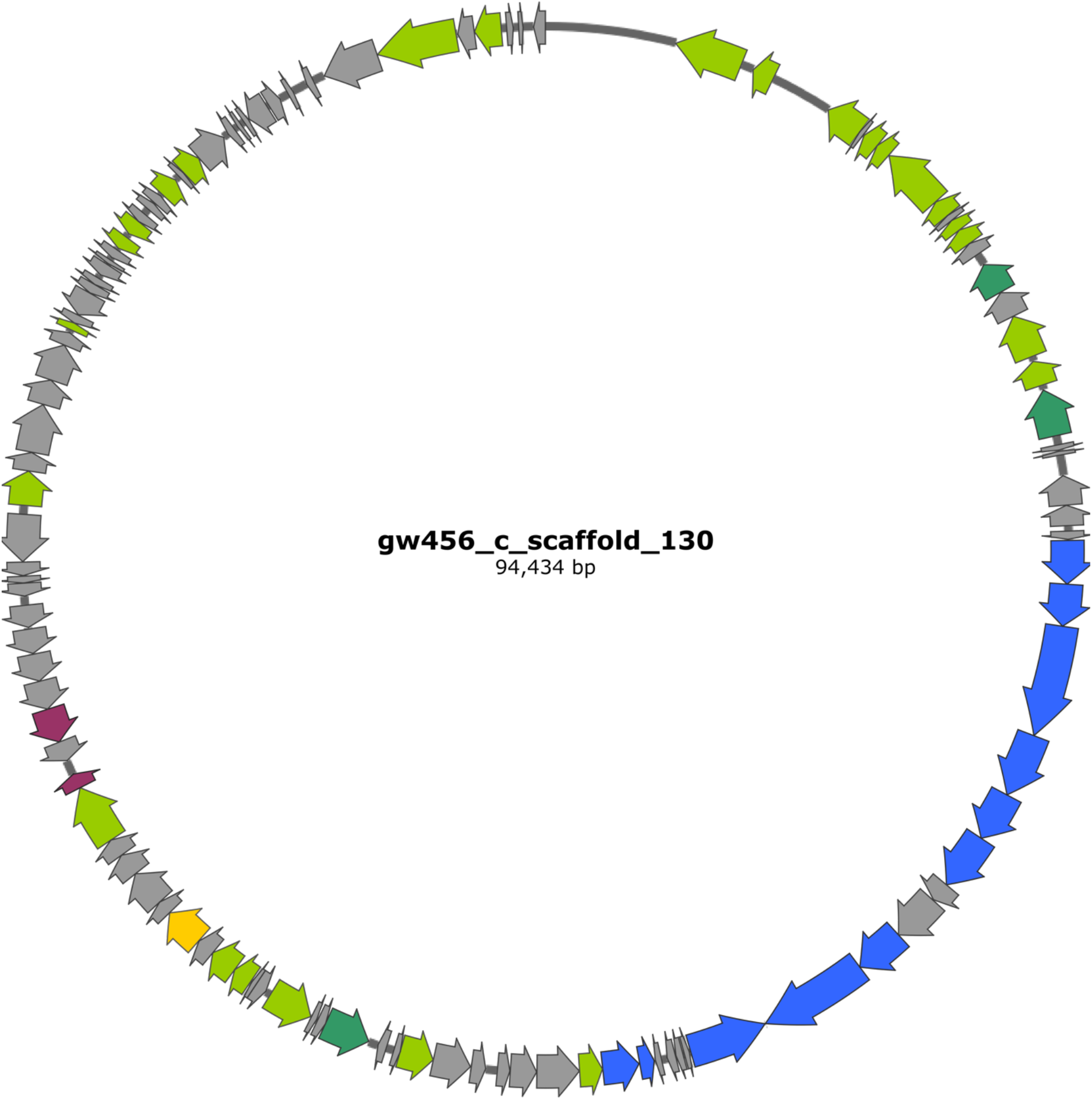
Example of a viral contig carrying auxiliary metabolite genes. Map of the virus (gw456_c_scaffold_130) from background groundwater with phage-related genes highlighted in green (darker green represents true hallmark genes of viruses), metal (copper, cobalt, zinc, cadmium, lead, mercury, arsenic) resistance genes highlighted blue, antibiotic (spectinomycin and fosfomycin) resistance genes highlighted in pink and metabolism (lactate dehydrogenase) gene in yellow. The viral contig was annotated via Prokka in Kbase^54^ and the annotation for virus-associated genes were updated on the map using virSorter^18^ predictions, details in Table S10.

## Conclusion

We demonstrate identification of novel viruses by leveraging plasmidome data for exploring environmental viral communities. Our analyses revealed the presence of novel viruses, likely representing new viral genera, in the underexplored groundwater environment. Using different datasets, we achieved bacterial host predictions for a substantial number of the viral sequences. Several of these phages encode genes related to signaling and tolerance mechanisms, thus likely augmenting ecosystem function by modifying the metabolism of their bacterial hosts. Interestingly, we find genes annotated to provide tolerance to metals, which is significant source of stress at this site. These predictions form the basis of future work on guiding phage isolation efforts and functional assessment of virus-host linkages. The ability to isolate phages would open new avenues for targeted manipulation of specific subsets of bacteria thus allowing for the systematic dissection of a microbiome for probing community dynamics and function.

## Acknowledgements

We thank Prof. Terry C. Hazen (University of Tennessee, Knoxville) for providing ORFRC groundwater samples for the isolation of bacterial strains. We thank Austin Hendricks and Jon Penterman for assistance as a part of the molecular biology MIT core facility; MIT BioMicroCenter (https://openwetware.org/wiki/BioMicroCenter). We thank Albert Tang, University of California, Berkeley for help with data analysis.

## Supplementary Information Figures

**Fig S1:**
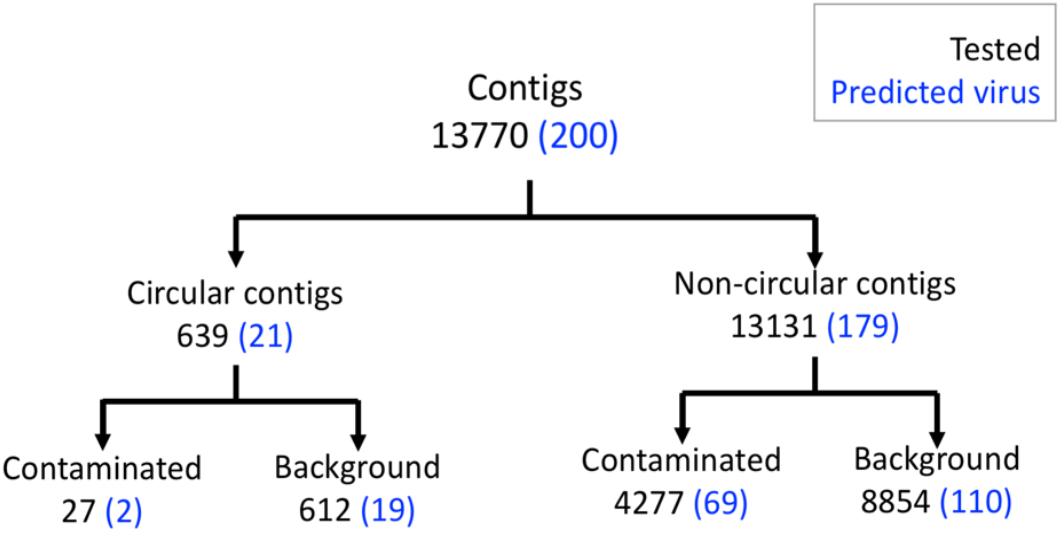
Breakdown of the all the contigs tested, with the those predicted to be viral highlighted in blue.

**Fig S2:**
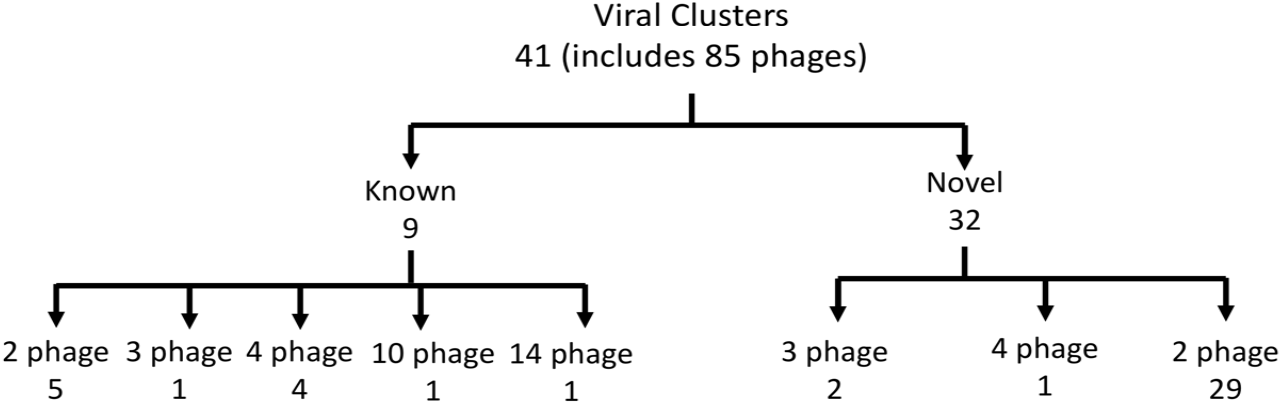
Distribution of viral clusters and the number of phages in each cluster.

**Fig S3:**
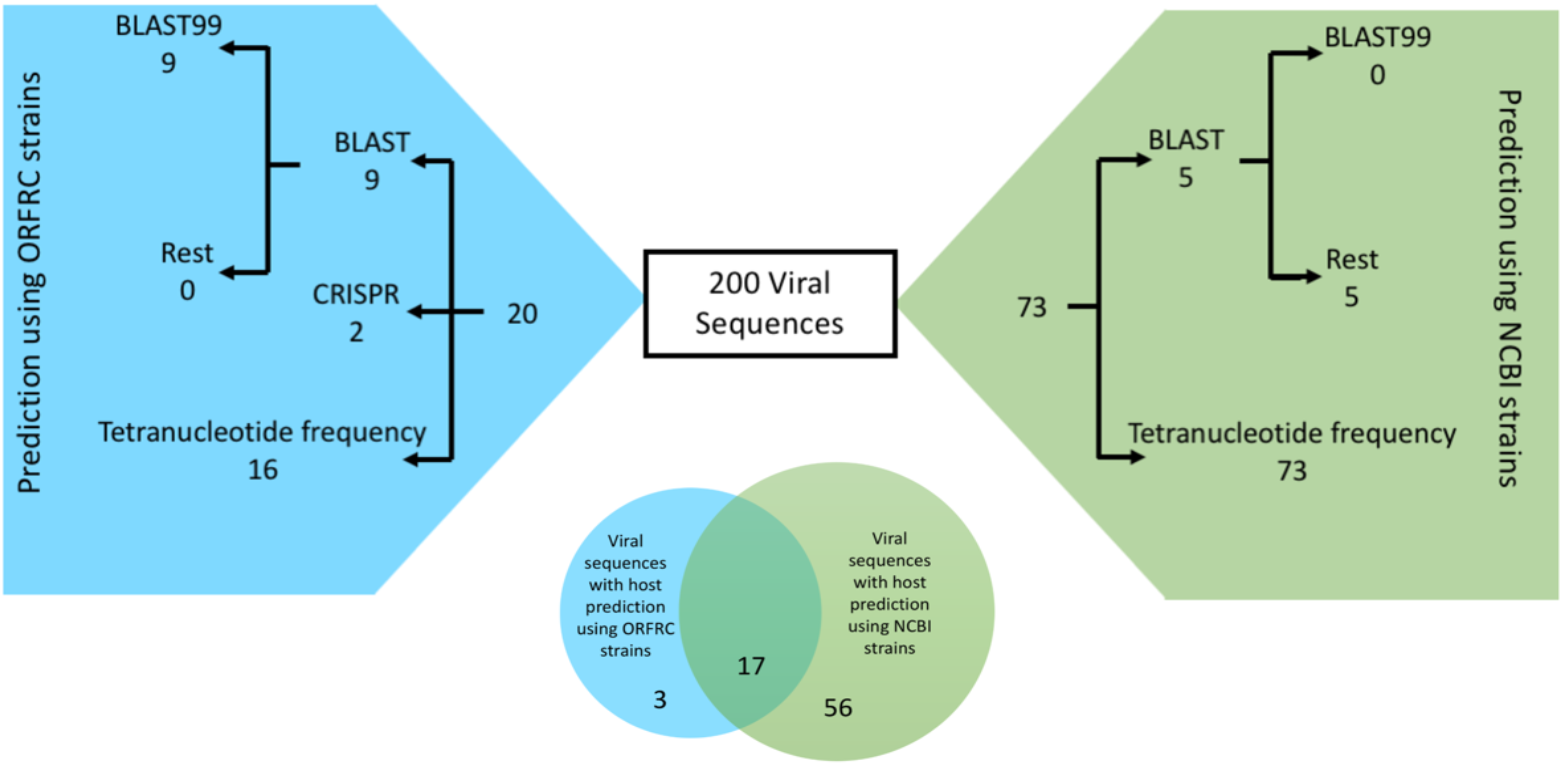
Details of numbers of viral sequences for which hosts are predicted using various bacterial host prediction methods on ORFRC and NCBI strains with whole genome sequences. Venn diagram shows the overlap of host prediction for viral sequences when using the ORFRC and NCBI strains.

Fig S4: Results of host association for virus GW456_nc_scaffold_4557. About 99.9% of the viral sequence (e-value = 0) could be found in 5 *Acidovorax* strains isolated from ORFRC.

**Fig S5:**
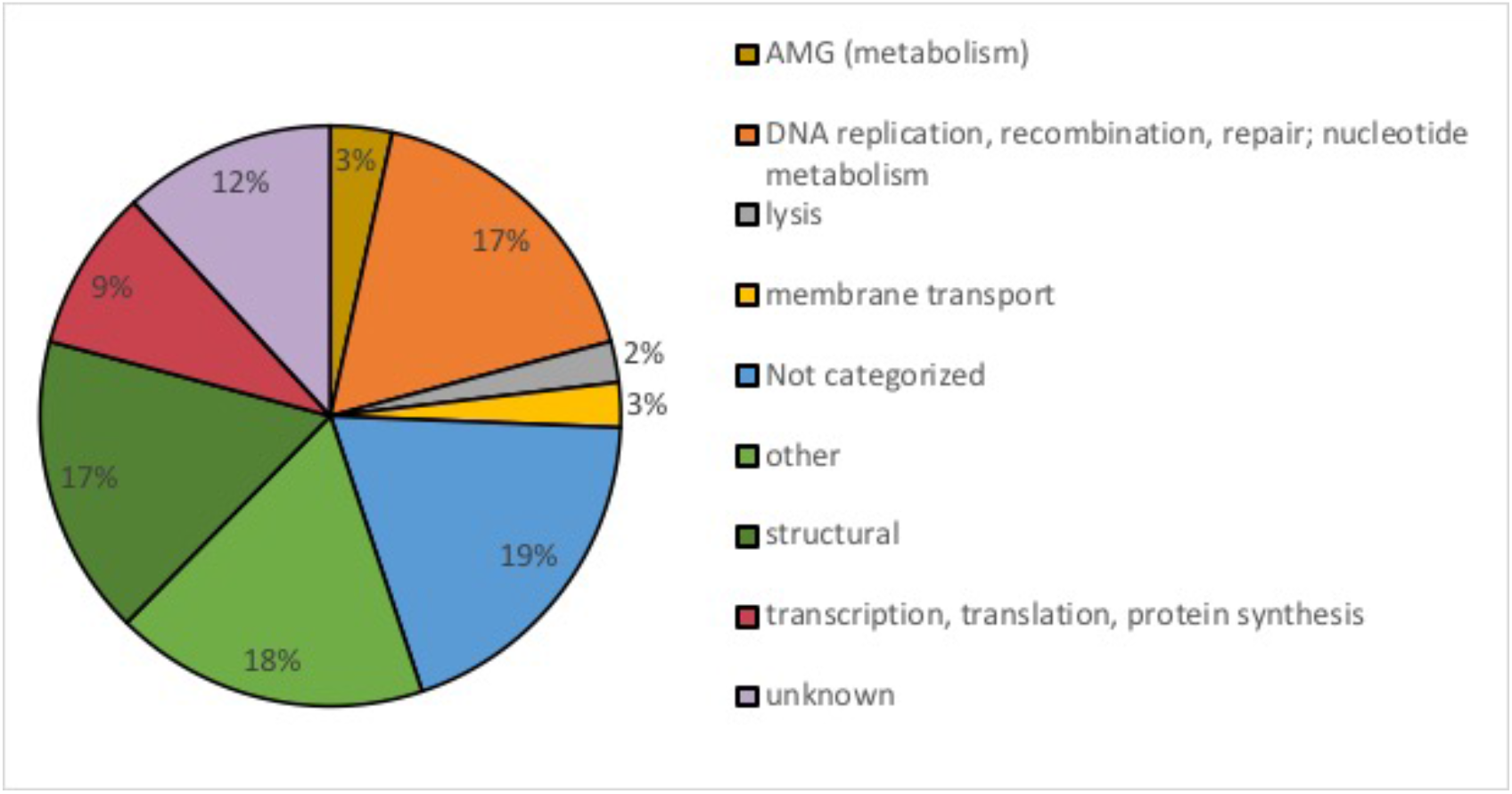
Major categories of 1486 virus encoded genes predicted via Pfam database. The hits were sorted into categories based on gene names as previously established based on AMGs seen in ocean viruses^34^.

## Supplementary Information Tables

Table S1: This file contains information on all contigs identified as viruses. These are sorted into categories phage (1, 2, 3) and prophage (4, 5, 6). Only the higher confidence categories 1, 2, 4 and 5 and considered as phage in this study.

Table S2: This file contains details of the 261 bacterial strains isolated from ORFRC.

Table S3: This file contains information on all 41 viral clusters that groundwater viruses fall into. This includes 32 novel clusters and 9 known clusters including known phages.

Table S4: Host association predictions based on ORFRC groundwater bacterial whole genome sequences.

Table S5: Host association predictions based on NCBI bacterial whole genome sequences.

Table S6: Breakdown of host assignments made using NCBI and ORFRC bacterial whole genome sequences.

Table S7: Comparison of host prediction between members of the same VC using NCBI and ORFRC bacterial whole genome sequences.

Table S8: Details of all the Pfam domains detected on the 200 viral sequences. Pfam domains related to metal resistance, antibiotic resistance and toxin-antitoxin encoding genes are also listed.

Table S9: Details of the 121 genes encoded on the viral sequence GW456_c_scaffold_130 including their location, size, directionality and description.

Table S10: Details of all 200 viral sequences with compilation of all analyses including virus size, viral cluster, host prediction, and AMG analysis.

## References

1. Sime-Ngando, T. Environmental bacteriophages: viruses of microbes in aquatic ecosystems. Front Microbiol 5, 355 (2014).

2. Brum, J.R. & Sullivan, M.B. Rising to the challenge: accelerated pace of discovery transforms marine virology. Nat Rev Microbiol 13, 147–159 (2015).

3. Rohwer, F. & Thurber, R.V. Viruses manipulate the marine environment. Nature 459, 207–212 (2009).

4. Breitbart, M., Bonnain, C., Malki, K. & Sawaya, N.A. Phage puppet masters of the marine microbial realm. Nat Microbiol 3, 754–766 (2018).

5. Coutinho, F.H., Gregoracci, G.B., Walter, J.M., Thompson, C.C. & Thompson, F.L. Metagenomics Sheds Light on the Ecology of Marine Microbes and Their Viruses. Trends Microbiol 26, 955–965 (2018).

6. Kauffman, K.M. et al. Viruses of the Nahant Collection, characterization of 251 marine Vibrionaceae viruses. Sci Data 5(2018).

7. Andreani, J., Verneau, J., Raoult, D., Levasseur, A. & La Scola, B. Deciphering viral presences: two novel partial giant viruses detected in marine metagenome and in a mine drainage metagenome. Virol J 15(2018).

8. Yu, D.T., Han, L.L., Zhang, L.M. & He, J.Z. Diversity and Distribution Characteristics of Viruses in Soils of a Marine-Terrestrial Ecotone in East China. Microb Ecol 75, 375–386 (2018).

9. Weynberg, K.D. Viruses in Marine Ecosystems: From Open Waters to Coral Reefs. Adv Virus Res 101, 1–38 (2018).

10. Appelo CA, P.D. Geochemistry, groundwater and pollution. CRC press (2004).

11. Watson, D., Kostka, J., Fields, M. & Jardine, P. The Oak Ridge field research center conceptual model. NABIR Field Research Center, Oak Ridge, TN (2004).

12. Bruce, G.M., Flack, S. M., Mongan, T. R. & Widner, T. E. Mercury releases from lithium enrichment at the Oak Ridge Y-12 plant: A reconstruction of historical releases and off-site doses and health risks. Reports of the Oak Ridge Dose Reconstruction (Tennessee Department of Health) (1999).

13. Rothschild, E.R., Turner, R.R., Stow, S.H., Bogle, M.A., Hyder, L.K., Sealand, O.M., & Wyrick, H.J. (1984).

14. Schulz, F. et al. Hidden diversity of soil giant viruses. Nat Commun 9(2018).

15. Ahlgren, N.A., Fuchsman, C.A., Rocap, G. & Fuhrman, J.A. Discovery of several novel, widespread, and ecologically distinct marine Thaumarchaeota viruses that encode amoC nitrification genes. The ISME journal, 1 (2018).

16. Edwards, R.A. & Rohwer, F. Viral metagenomics. Nat Rev Microbiol 3, 504–510 (2005).

17. Kothari, A. et al. Large Circular Plasmids from Groundwater Plasmidomes Span Multiple Incompatibility Groups and Are Enriched in Multimetal Resistance Genes. MBio 10(2019).

18. Roux, S., Enault, F., Hurwitz, B.L. & Sullivan, M.B. VirSorter: mining viral signal from microbial genomic data. Peerj 3(2015).

19. Bolduc, B., Youens-Clark, K., Roux, S., Hurwitz, B.L. & Sullivan, M.B. iVirus: facilitating new insights in viral ecology with software and community data sets imbedded in a cyberinfrastructure. Isme Journal 11, 7–14 (2017).

20. Bolduc, B. et al. vConTACT: an iVirus tool to classify double-stranded DNA viruses that infect Archaea and Bacteria. Peerj 5(2017).

21. Shannon, P. et al. Cytoscape: A software environment for integrated models of biomolecular interaction networks. Genome Res 13, 2498–2504 (2003).

22. Parks, D.H., Imelfort, M., Skennerton, C.T., Hugenholtz, P. & Tyson, G.W. CheckM: assessing the quality of microbial genomes recovered from isolates, single cells, and metagenomes. Genome Res 25, 1043–1055 (2015).

23. Martin, M. Cutadapt removes adapter sequences from high-throughput sequencing reads. EMBnet. journal 17, 10–12 (2011).

24. Bolger, A.M., Lohse, M. & Usadel, B. Trimmomatic: a flexible trimmer for Illumina sequence data. Bioinformatics 30, 2114–2120 (2014).

25. Bankevich, A. et al. SPAdes: A New Genome Assembly Algorithm and Its Applications to Single-Cell Sequencing. J Comput Biol 19, 455–477 (2012).

26. Edgar, R. SINTAX: a simple non-Bayesian taxonomy classifier for 16S and ITS sequences. BioRxiv, 074161 (2016).

27. Lagesen, K. et al. RNAmmer: consistent and rapid annotation of ribosomal RNA genes. Nucleic Acids Res 35, 3100–3108 (2007).

28. Cole, J.R. et al. Ribosomal Database Project: data and tools for high throughput rRNA analysis. Nucleic Acids Res 42, D633–D642 (2014).

29. Roux, S. et al. Minimum Information about an Uncultivated Virus Genome (MIUViG). Nat Biotechnol 37, 29–37 (2019).

30. Edwards, R.A., McNair, K., Faust, K., Raes, J. & Dutilh, B.E. Computational approaches to predict bacteriophage-host relationships. Fems Microbiology Reviews 40, 258–272 (2016).

31. Andersson, A.F. & Banfield, J.F. Virus population dynamics and acquired virus resistance in natural microbial communities. Science 320, 1047–1050 (2008).

32. Mizuno, C.M., Rodriguez-Valera, F., Kimes, N.E. & Ghai, R. Expanding the Marine Virosphere Using Metagenomics. Plos Genet 9(2013).

33. Roux, S., Hallam, S.J., Woyke, T. & Sullivan, M.B. Viral dark matter and virus-host interactions resolved from publicly available microbial genomes. Elife 4(2015).

34. Roux, S. et al. Ecogenomics and potential biogeochemical impacts of globally abundant ocean viruses. Nature 537, 689−+ (2016).

35. Bland, C. et al. CRISPR Recognition Tool (CRT): a tool for automatic detection of clustered regularly interspaced palindromic repeats. Bmc Bioinformatics 8(2007).

36. Biswas, A., Gagnon, J.N., Brouns, S.J.J., Fineran, P.C. & Brown, C.M. CRISPRTarget: Bioinformatic prediction and analysis of crRNA targets. Rna Biol 10, 817–827 (2013).

37. Ogilvie, L.A. et al. Genome signature-based dissection of human gut metagenomes to extract subliminal viral sequences. Nat Commun 4(2013).

38. Marcais, G. & Kingsford, C. A fast, lock-free approach for efficient parallel counting of occurrences of k-mers. Bioinformatics 27, 764–770 (2011).

39. Edgar, R.C. MUSCLE: a multiple sequence alignment method with reduced time and space complexity. Bmc Bioinformatics 5, 1–19 (2004).

40. Kumar, S., Stecher, G. & Tamura, K. MEGA7: Molecular Evolutionary Genetics Analysis Version 7.0 for Bigger Datasets. Mol Biol Evol 33, 1870–1874 (2016).

41. El-Gebali, S. et al. The Pfam protein families database in 2019. Nucleic Acids Res 47, D427–D432 (2019).

42. Eddy, S.R. Accelerated Profile HMM Searches. Plos Comput Biol 7(2011).

43. Gorlas, A., Krupovic, M., Forterre, P. & Geslin, C. Living Side by Side with a Virus: Characterization of Two Novel Plasmids from Thermococcus prieurii, a Host for the Spindle-Shaped Virus TPV1. Appl Environ Microb 79, 3822–3828 (2013).

44. Arnold, H.P. et al. The genetic element pSSVx of the extremely thermophilic crenarchaeon Sulfolobus is a hybrid between a plasmid and a virus. Mol Microbiol 34, 217–226 (1999).

45. Lima-Mendez, G., Van Helden, J., Toussaint, A. & Leplae, R. Reticulate representation of evolutionary and functional relationships between phage genomes. Mol Biol Evol 25, 762–777 (2008).

46. Ge, X.X. et al. Iron- and aluminium-induced depletion of molybdenum in acidic environments impedes the nitrogen cycle. Environmental Microbiology 21, 152–163 (2019).

47. Waldor, M.K. & Mekalanos, J.J. Lysogenic conversion by a filamentous phage encoding cholera toxin. Science 272, 1910–1914 (1996).

48. Breitbart, M., Thompson, L.R., Suttle, C.A. & Sullivan, M.B. Exploring the Vast Diversity of Marine Viruses. Oceanography 20, 135–139 (2007).

49. Yooseph, S. et al. The Sorcerer II Global Ocean Sampling expedition: Expanding the universe of protein families. Plos Biol 5, 432–466 (2007).

50. Hemme, C.L. et al. Metagenomic insights into evolution of a heavy metal-contaminated groundwater microbial community. Isme Journal 4, 660–672 (2010).

51. Hemme, C.L. et al. Lateral gene transfer in a heavy metal-contaminated-groundwater microbial community. MBio 7, e02234–02215 (2016).

